# PreDigs: a Database of Context-specific Cell-type Markers and Precise cell subtypes for Digestive Cell Annotation

**DOI:** 10.1101/2024.10.29.620788

**Authors:** Jiayue Meng, Mengyao Han, Yuwei Huang, Liang Li, Yuanhu Ju, Daqing Lv, Xiaoyi Chen, Liyun Yuan, Guoqing Zhang

**Author notes:** Corresponding author(s). (Guoqing Zhang), n (Liyun Yuan). Equal contribution (Jiayue Meng, Mengyao Han, Yuwei Huang).

## Abstract

Research on cell type markers aids investigators in exploring the diverse cellular compositions within gastrointestinal tumors, enhancing our understanding of tumor heterogeneity and its implications for disease progression and treatment response. However, issues such as the integration of large-scale datasets and the lack of standardized cell type identification hinder comprehensive characterization. Here, we developed a user-friendly web interface called PreDigs (Predicted Signatures in Digestive System), which offers 124 tailored scRNA-seq datasets available for download, encompassing over 3.4 million cells. After unsupervised clustering, we unified the identification and naming of subtype labels, ultimately constructing a cell ontology tree that includes 142 cell types, with up to eight hierarchical levels. Meanwhile, we calculated three different context-specific cell-type markers—’Cell Markers’, ‘Subtype Markers’, and ‘TPN Markers’—based on various application requirements within or across tissues. Through the integrated analysis of PreDigs gastrointestinal data, we identified distinct cell subpopulations exclusive to tumors, one of which corresponds to tumor-specific endothelial cells (TEC). Furthermore, PreDigs offers online cell annotation tools that empower users to perform single-cell classification with greater flexibility, accessible at https://www.biosino.org/predigs/.

## Introduction

Digestive cancers, including esophageal, gastric, liver, pancreatic and colorectal cancers, account for more than 50% of global cancer-related morbidity and mortality(1). As solid tumors, the complex cellular components with heterogeneous cancer, tissue-resident and tumor-infiltrating cells in tumor micro-environment (TME) influence tumor cell growth and development(2, 3). Leveraging massive single-cell RNA sequencing (scRNA-seq) data has given us the opportunity for the comprehensive profiling of gene expression at the single-cell level, enabling researchers to uncover cellular composition diversity(4), distinguish cell lineages(5), characterize TME for the digestive cancers(6). Precise cell type classification is an important step in the single-cell analysis for cancer research and is also the key to understand intra-tumor heterogeneity (7-10).

Until now, great effects have been made to develop several cell-type markers data portals and sources for single-cell classification(11-20). Among them, notable databases include Cell ontology(11), CellMarkers(12, 13), PanglaoDB(15), Cell Taxonomy(19) and CellSTAR(20). Cell Ontology (CL), a widely-used framework for representing cell types, establishes hierarchical relationships for approximately 2,600 cell types, although most of these types are not associated with assessable cell markers. CellMarker, the first database to compile a curated collection of 27,166 cell markers for 3,123 cell types in human and mouse, does not incorporate single-cell transcriptome profiles for these markers. In contrast, databases like PanglaoDB have compiled cell types, markers and scRNA-seq datasets, but inconsistencies in dataset preprocessing procedures and the absence of a unified nomenclature for cell types, such as the adoption of the CL standardization, hinder direct cross-dataset comparisons. Cell Taxonomy and CellSTAR have addressed these issues, yet they do not support cross-tissue comparisons, such as the gene expression differences of the same cell type in normal and tumor tissues. This gap restricts our understanding of cell-type markers in the TME and the precise annotation of TME cell types (37).

Here, we present PreDigs (Predicted Signatures in Digestive System), a comprehensive database that integrates extensive scRNA-seq datasets and utilize sophisticated reference data for cell subtype annotation to construct a detailed cell ontology tree encompassing 142 distinct cell types with up to eight layers, allowing each node within the hierarchy to be linked to relevant expression profiles, thereby enabling comparative analyses across diverse tissues and datasets. Furthermore, PreDigs affords three types of markers tailored for different research contexts: direct comparison of DEG profiles for a cell type across tissue types (‘Cell Markers’), delineation of gene expression differences within cell subtypes via cross-tissue single-cell analysis (‘Subtype Markers’), and identification of cell types specifically distributed within tumor tissue and their gene functions in KEGG pathways (‘TPN Markers’). The PreDigs interface allows for easy retrieval of cell-type markers and scRNA-seq datasets, supporting in-depth analysis of gene expression and functional enrichment. Additionally, an online annotation tool is provided, enhancing single-cell classification efficiency and accuracy by enabling annotation based on both classical markers and tailored scRNA-seq reference datasets.

## Results

### Overview of data management and integration in PreDigs

In summary, we obtained 124 fine-processed scRNA-seq datasets of adult digestive tissues (see Methods and Figure S1). These datasets include three distinct types of digestive tissues (normal tissue, paracancerous tissue, and tumor tissue) across five digestive organs (intestine, pancreas, esophagus, liver, and stomach), and approximately 3.4 million single cells (Figure 1A). Besides, to structurally describe digestive cell types, we systematically developed a cell ontology tree of digestive cells with up to eight levels, incorporating 142 cell types, by referencing the Cell Ontology (CL) database (see Methods and Figure S1). Among them, 74% (105 types) correspond to standardized CL IDs within the CL database. The first layer of the cell ontology tree is divided into eleven groups (Figure 1B), with the four main types being immune cells, stromal cells, secretory cells, and epithelial cells. For instance, in Figure 1B, immune cells are categorized into two main lineages—lymphocyte and myeloid leukocyte—and further delineated into 64 hierarchically distinct subtypes. Furthermore, PreDigs has enhanced the annotation of the cell ontology tree by incorporating 3,693 canonical markers across 74 cell types derived from a comprehensive review curation, thereby enriching the associated cell-type marker information (see Methods).

**Figure 1.**
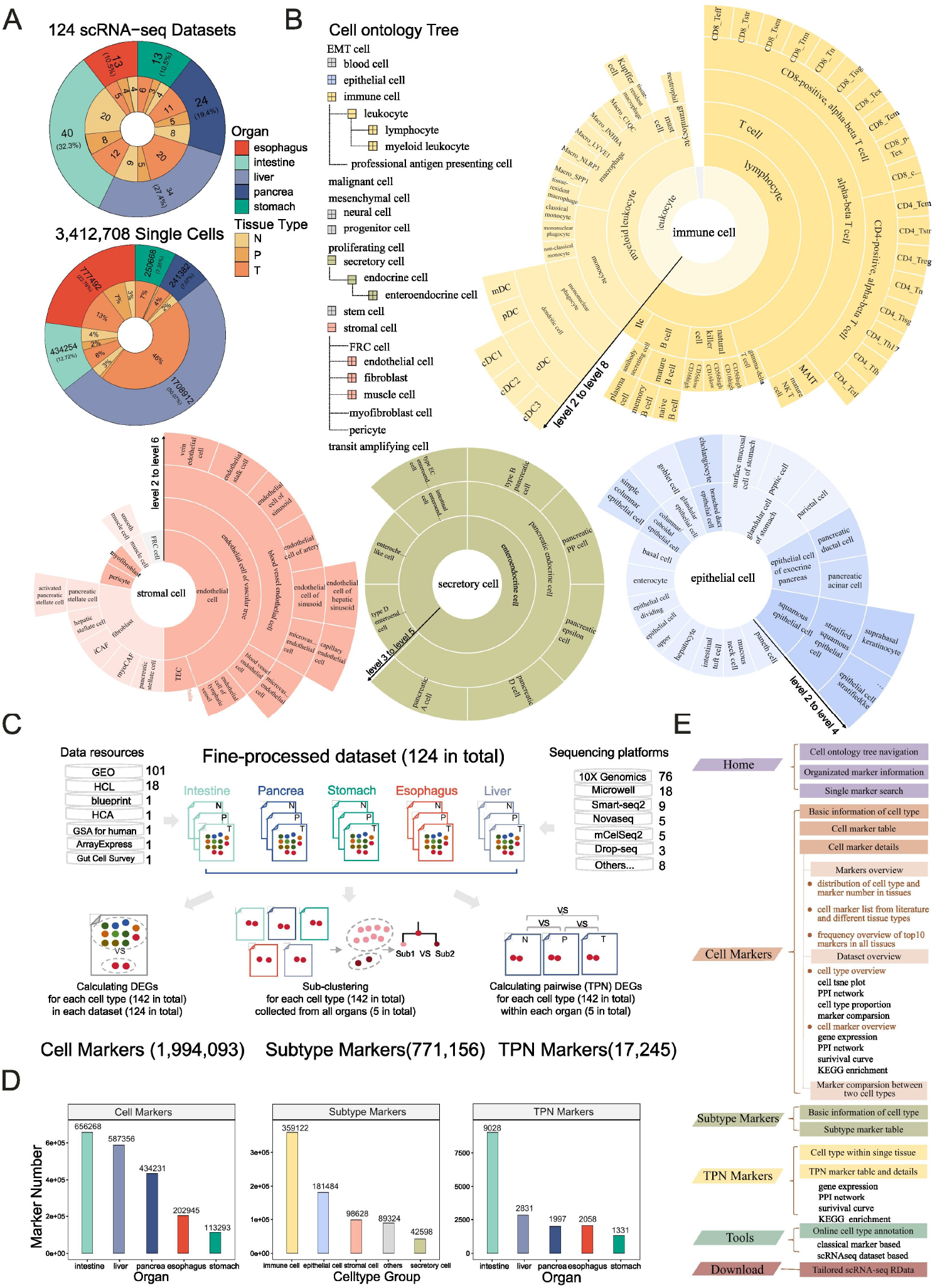
The summary of PreDigs data. A. The pie plots show dataset and single-cell number across different digestive organ and tissue types, respectively. B. An illustration of the cell ontology tree with 8 levels. C. Overview of data processing workflow for context-specific markers in PreDigs. D. The statistics on the number of three types of context-specific markers across different digestive tissues. E. Schematic overview of the PreDigs database interface.

After obtaining the tailored scRNA-seq datasets and cellular ontology tree, we employed different strategies to calculate three context-specific types of cell markers to suit various application scenarios (see Methods and Figure 1C). Firstly, within each dataset, we preliminarily screened DEGs with significantly higher expression in specific cell types compared to others (‘Cell Markers’). We then compared these DEG sets of the same cell type across different tissue types to clarify their commonalities and differences. Secondly, we utilized integrative analysis of cross-tissue single-cell data to thoroughly investigate gene expression differences among various subtypes within over 140 cell types (‘Subtype Markers’). Finally, we focused on multi-dataset integration within specific tissue contexts to systematically assess the expression differences of specific cell types across three representative tissue types and conducted corresponding functional enrichment analyses (‘TPN Markers’).

Through the above workflows, we obtained a total of 2,782,494 marker records (Figure 1D), including 1,994,093 ‘Cell Markers’, 17,245 ‘TPN Markers’, and 771,156 ‘Subtype Markers’. We found that the number of ‘Cell Markers’ in the intestine, liver, and pancreas significantly increased compared to the number of those in the esophagus and stomach. Furthermore, this increasing trend aligns with the quantity of datasets. However, the number of ‘TPN Markers’ in intestinal tissues is significantly higher than in other gastrointestinal tissues. This may be due to the greater variability in TPN among the intestinal samples compared to other tissues (Figure 1D).

Here, we developed PreDigs, a web resource for viewing cell-type markers base on different contextual needs and facilitating browsing and comparison by organ, tissue, and cell type, as well as downloading tailored scRNA-seq datasets. It includes six key pages (Figure 1E), with the homepage offering a cell ontology tree, data summary and marker preparation details. On the following three marker pages, we provide a marker table with extensive links to single gene expression and KEGG pathway enrichment results. This setup enables users to delve into gene expression features and functional implications upon obtaining the marker list. Finally, the integration of these data drives an online tool designed for accurate annotation of digestive cells.

### Interface for browsing three types of cell markers

In PreDigs, a cell ontology tree is always provided on the left side of all web pages, allowing users to browse and select cell types (Figure 2). Three context-specific markers, i.e. ‘Cell Markers’ (C), ‘Subtype Markers’ (S), and ‘TPN Markers’ (T) were respectively displayed across organs, tissues, and cell types. Once a cell type is selected on the left cell ontology tree, detailed information including name, ontology ID, and description, will be displayed first. Meanwhile, the marker list will be shown on the following table (Figure 2A). Details for each marker gene can be accessed by clicking the eye icon on the right (Figure 2B-D). Here, we use “Cell Markers” page as an example to provide a detailed introduction to browsing marker information, while the content on the other pages is relatively simpler and similar.

**Figure 2.**
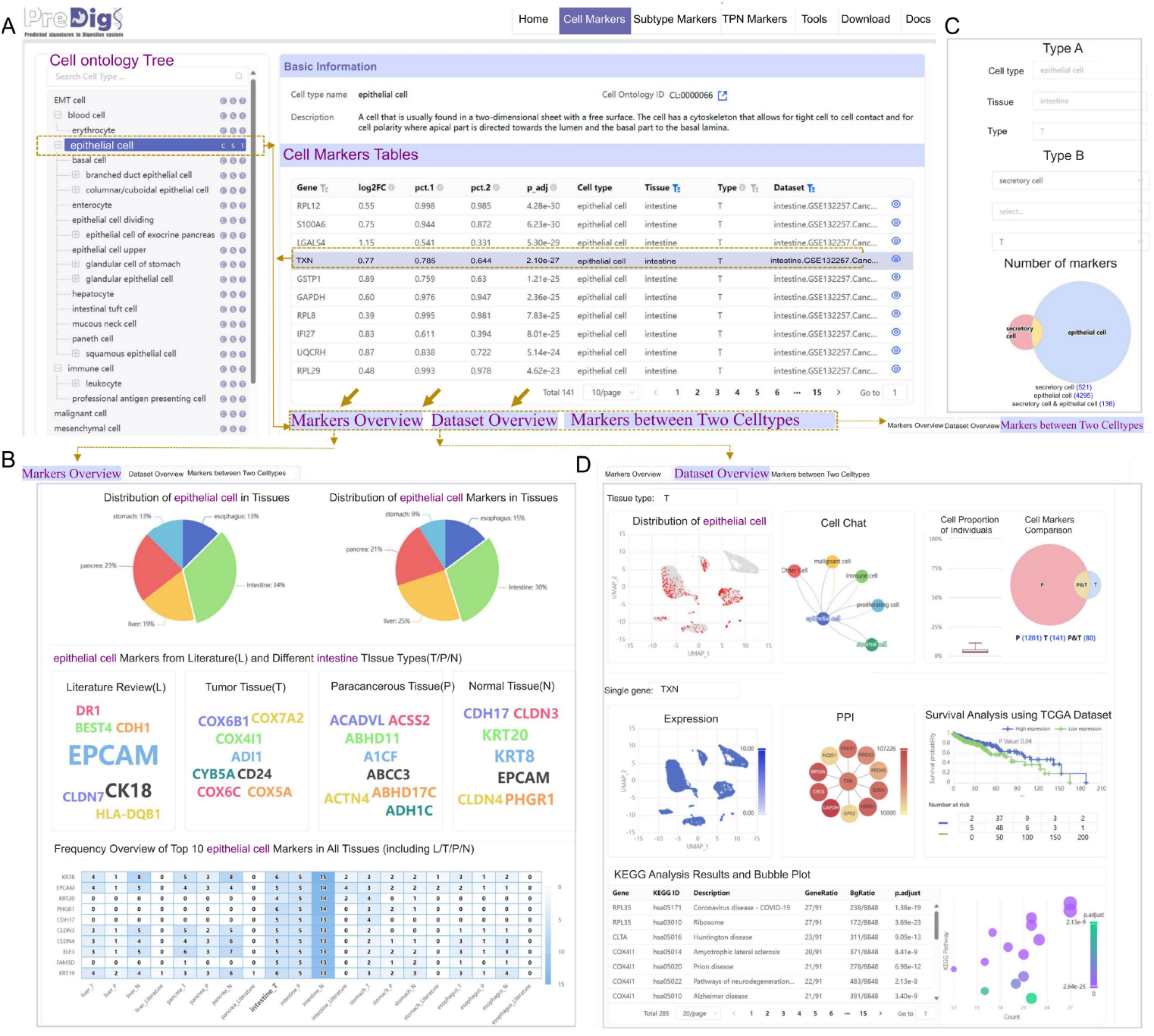
Screenshots of “Cell Markers” interface in PreDigs. A. “Cell Markers” interface includes a cell ontology tree, markers table and three tabs for different functions. B. ‘Markers Overview’ tab for cell marker statistics. C. ‘Markers between Two Celltypes’ tab for comparative analysis. D. ‘Dataset Overview’ tab for expression profiles in scRNA-seq datasets.

*‘Markers overview’* tab on “*Cell Marker*” page: as shown in Figure2B, the tab presents an overview of the “Cell Markers” contained within the user-selected cell type, including the quantitative distribution of these markers across gastrointestinal tissues and comparisons among different tissue types. Here, we also counted the occurrences of markers in each tissue and visualized the top 10 markers with the highest frequencies, enabling users to gain a deeper understanding of these genes’ expression characteristics in different tissues.

*‘Markers between Two Celltypes’* tab on “*Cell Marker*” page: as shown in Figure2C, this online cell-type comparison tab helps researchers to quickly contrast the marker profiles of two distinct cell types within a particular tissue. Users can select the cell types and instantly view the similarities and differences in their marker lists.

*‘Dataset Overview’* tab on “*Cell Marker*” page: as shown in Figure2D, this tab focuses on the current single dataset, allowing users to explore the clustering relationships (tSNE) between the selected cell type and other cell types, intercellular communication (Cell Chat), differences in cell markers (Venn), as well as the expression levels of selected genes (tSNE), interactions with other genes (PPI), and prognostic impact (survival analysis), functional analysis (KEGG).

“Subtype Marker” page provides a research platform for in-depth analysis of subtypes within a given cell type on the left cell ontology tree (Figure S2). We performed more refined clustering of each cell type, and users can explore the distribution of subtypes across organs and tissue types. This helps users to gain a better understanding of diversity and heterogeneity of cell subtypes across digestive tissues to identify specific subtype. Additionally, users can also browse detailed markers of each subtype on the following table.

“TPN Markers” page focuses on examining DEGs of the same cell type in pairwise comparison across distinct tissue types, i.e. tumor (T), paracancerous (P), and normal (N) tissues. It will display the distribution of the selected cell type in the organ, as well as in each individual sample (Figure S3). All pairwise comparison markers will also be shown on the following table. Similarly, details for each TPN marker gene can be accessed by clicking the eye icon on the right. Detailed information including gene pression (UMAP), gene network (PPI), functional analysis (survival analysis and KEGG enrichment) was all displayed. These analytical outcomes aid researchers in targeting signaling pathways and understanding their biological significance for subsequent studies.

### Online cell type annotation platform

PreDigs also provides an interactive platform with two methods: cell markers and tailored reference datasets, enabling users to perform cell type annotation on the “Tool” page (Figure 3). The platform accommodates the upload of single-cell expression matrices and metadata tables that users have previously queried. Upon uploading, users can select from a variety of parameters, including automated annotation tools, multiple reference data, and options for customizing the annotation process. For cell marker annotation, users are provided with a comprehensive list of marker genes for all cell types associated with the selected tissue, with the freedom to exclude any unnecessary types. For tailored reference datasets, users can choose one or more reference datasets to serve as the reference data. After submitting their selections, users will receive tSNE plots displaying cell group names and prediction labels, along with downloadable files containing the predicted labels and customized feature gene lists, enhancing the accessibility and efficiency of working with the PreDigs database.

**Figure 3.**
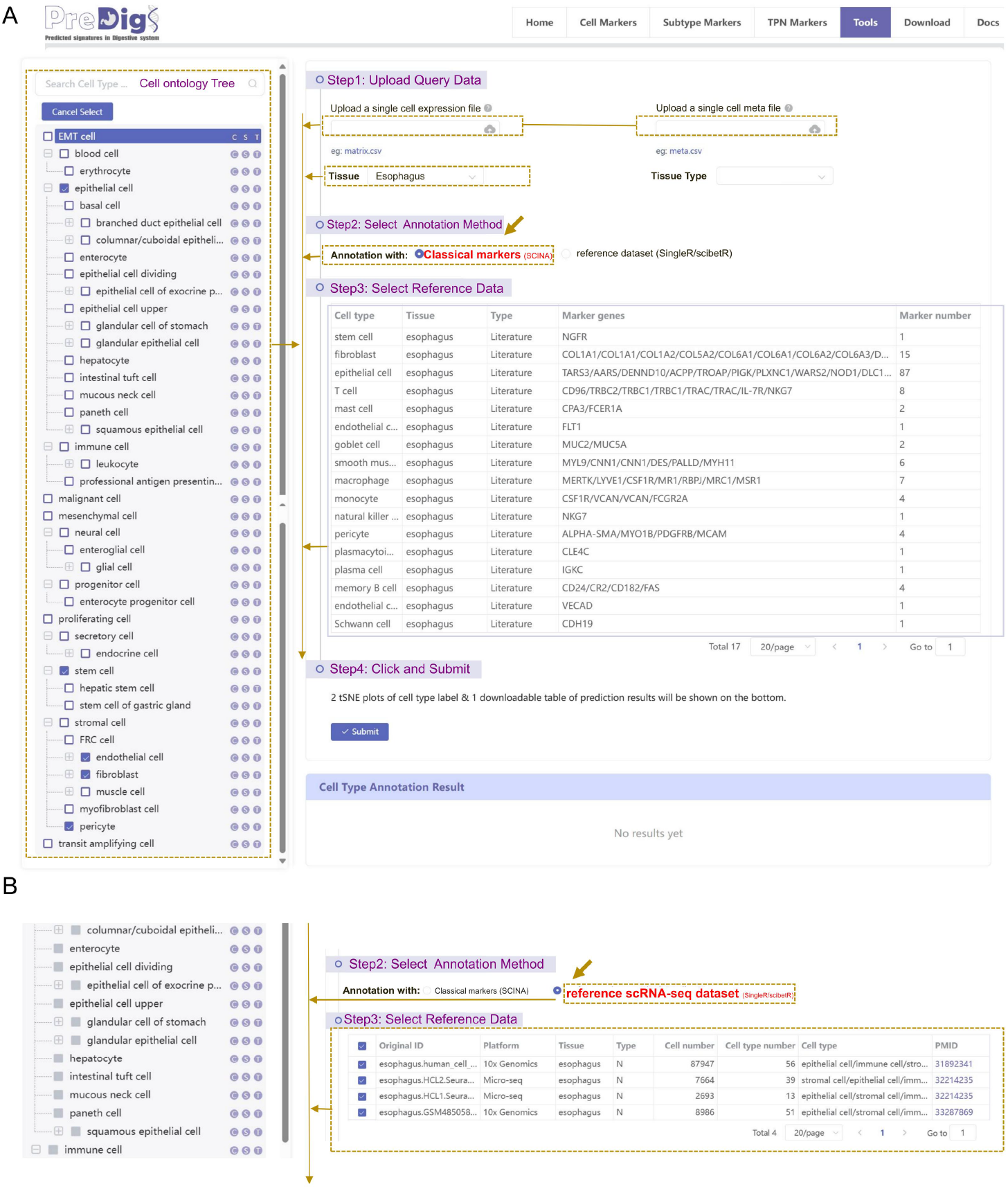
Screenshots of “Tools” interface in PreDigs. A. Step-by-step instructions for annotation using classical markers as reference data. B. Step-by-step instructions for annotation utilizing single or multiple scRNA-seq datasets as reference data.

### Utility of PreDigs reference for revealing cell type heterogeneity

In this study, we first used endothelial cell as an example and performed the large-scale integration of 124 scRNA-seq datasets from 5 digestive organs in the PreDigs database. This integration enabled us to categorize 96,429 cells into 8 distinct subtypes (Figure 4A). Notably, we identified a subtype comprising 20,019 single cells, which was predominantly found in tumor tissues. We designated these cells as gastrointestinal tumor-specific endothelial (TEC) cells. Furthermore, we further identified 375 ‘Subtype’ markers specifically highly expressed in TECs compared to other endothelial subtypes. KEGG functional enrichment results showed that half of the top 10 enriched pathways for these markers were related to cancer, including “Pathway in Cancer”, “PI3K-Akt signaling pathway”, “Small cell lung cancer” etc., effectively depicting the pathological state of endothelial cells within tumor tissue (Figure 4B).

**Figure 4.**
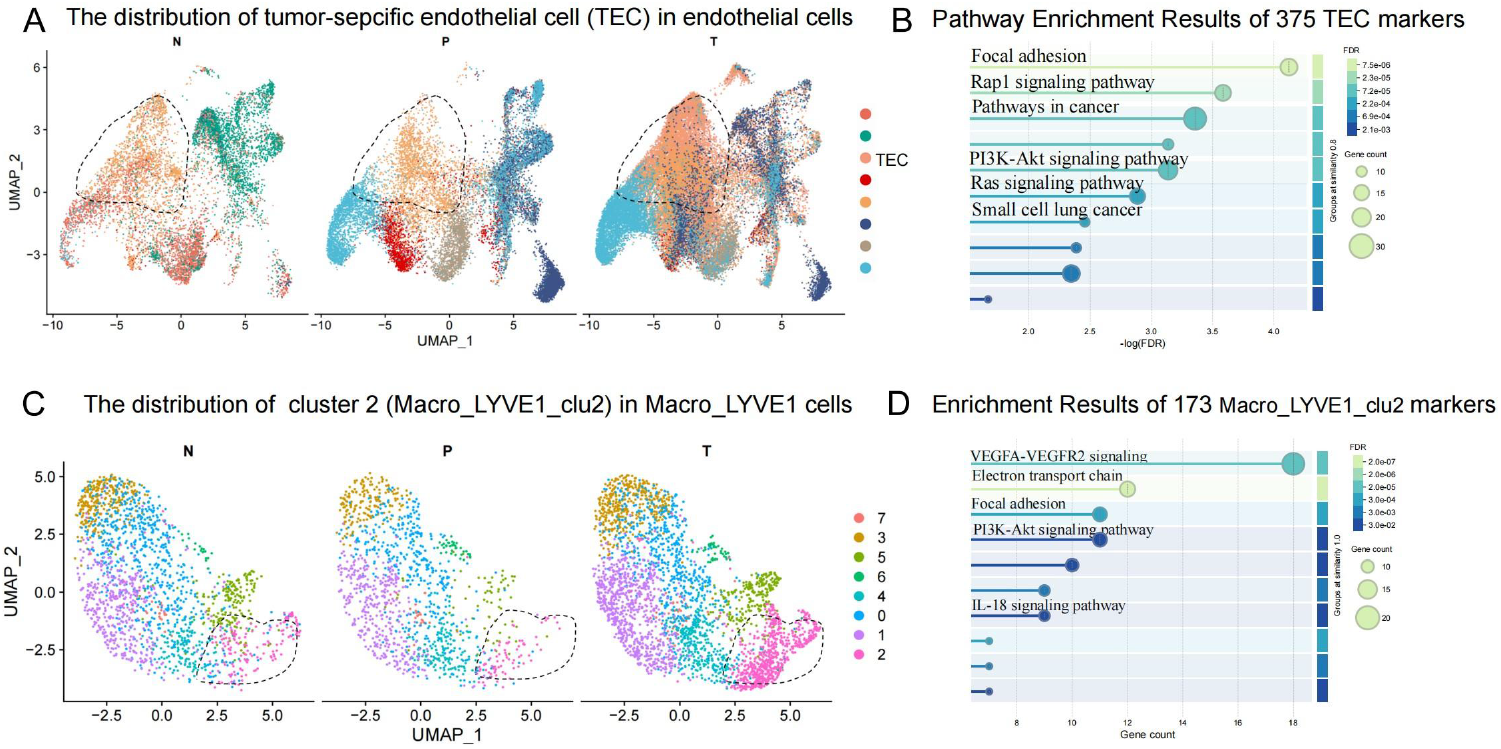
The utility of PreDigs reference for cell subtype annotation. A. Unsupervised clustering results of endothelial cells from the cross-tissue integration of PreDigs data. B. KEGG enrichment results of 375 ‘Subtype’ markers which are up-regulated in TEC compared to other endothelial subtypes. C. Unsupervised clustering results of Macro_LYVE1 cells resulting from the cross-tissue integration of PreDigs data. D. Wiki pathway enrichment results for the 173 ‘Subtype’ markers which are up-regulated in cluster 2 relative to other subclusters within the Macro_LYVE1 cell population.

Following a similar approach, we integrated and clustered 6,346 Macro_LYVE1 cells into eight subclusters, and we found that cluster2 (‘Macro_LYVE1_clu2’) was exclusively present in tumor tissues (Figure 4C). We then identified 173 ‘Subtype’ markers that were highly expressed in this cluster relative to other Macro_LYVE1 subclusters, which exhibited significant enrichment in the “IL-18 signaling pathway”, awaiting further biological investigation (Figure 4D).

## Discussion

We believe one of the key strengths of PreDigs lies in its comprehensive coverage of digestive tissue types and the standardized cell ontology tree, offering unique features compared to existing resources. Each cell type node is accompanied by thorough marker calculations, which we hope will make single-cell annotation more accessible across various research contexts, especially for cross-tissue comparisons to explore cellular heterogeneity and similarities among digestive organs. However, we acknowledge that the construction of this cell ontology tree currently relies heavily on available marker annotations, and we look forward to incorporating further experimental validations to improve accuracy.

The initial version of PreDigs focuses on gastrointestinal cancers, partly due to the complexity of constructing a new cell-type hierarchy and the extensive effort required for large-scale data integration. Looking ahead, we hope to expand PreDigs to include additional cell and tissue types, particularly by integrating datasets for digestive tumor subtypes not yet represented.

As multi-omics data integration methods advance, including spatial transcriptomics and single-cell TCR sequencing, we anticipate that these approaches will provide valuable insights into the tumor microenvironment, enhancing the database’s relevance. We also aim to incorporate large language models to further support single-cell classification, with the hope that PreDigs will continue to grow as a useful tool for in-depth research.

## Method

### Dataset collection and pre-processing

The 99 original scRNA-seq datasets of the adult digestive system were collected from seven main public data resources, including NCBI Gene Expression Omnibus (GEO), Human Cell Landscape (HCL)(21), Human cell Atlas (HCA), ArrayExpress, GSA for Human, Gut Cell Survey and blueprint. These datasets originate from various sequencing platforms, including 10X Genomics, Microwell, Smart-seq2, Drop-seq, STRT-seq, Novaseq, Fluidigm and others. The gene expression matrices were downloaded, as well as the meta-data tables containing cell type labels (if accessible in publications). Through manually reviewing each literature and supplemental materials, we curated the meta-information of each dataset, including participant, cell number, cell type number, cancer type, technology and dataset sources. Curated meta-information was elaborated in the database.

Before standardized pre-processing of 99 diverse datasets, we subdivided these into 124 independent data subsets categorized by three tissue types: N (normal tissue), P (paracancerous tissue), and T (tumor tissue). All the scRNA-seq data were processed using R (v3.6.1) with uniform methods except special explanation. Doublets removal was performed by DoubletFinder v2.2.0 (22)(7% per 10000 cells) for all datasets. Seurat v2.3.2 (23) was employed to perform a series of analysis: quality control (200 < nfeature < 5000 and MT percentage < 10%), normalization, Principal Component Analysis (PCA), dimension reduction, cell clustering with top 30 principal component numbers and UMAP (Uniform Manifold Approximation and Projection) visualization and differential gene expression (DGE) analysis through “FindAllMarkers” function with wilcoxon rank sum test (adjusted P value < 0.000005, log2 fold change > 1.5). Full-length sequencing generated datasets were mainly normalized by Transcript Per Million method instead of Seurat workflow.

Batch removal and data integration were performed by Harmony v1.0 (24) and canonical correlation analysis of Seurat. To integrate the data, we first identified variable genes in each dataset with the “FindVariableFeatures” function. We then scaled the data using the “ScaleData” function and proceeded to integrate different datasets with Seurat’s “FindIntegrationAnchors” and “IntegrateData” functions, with parameters set as follows: k.filter=50, resolution=0.5, and dims=30. Online analytics platform ViPMAP was assisted in above processes (https://www.biosino.org/vipmap/).

### Cell type annotation and clustering

We performed unified cell type annotations on the preprocessed gene expression matrix. The annotation process for healthy gastrointestinal tissues mainly consisted of the following three steps: First, we used the tool CellAssign v0.99.2(25) for automated cell type prediction; next, manual annotation was conducted based on the expression of typical cell markers; then, by comparing the re-annotated results with the original literature to determine cell labels; finally, the cell label names were manually curated based on standardized terms of the Cell Ontology database. For malignant cells classification in tumor tissues, we additionally fine-tuned the annotation by combining the expression of typical tumor cell marker genes (EPCAM, TP53, etc.).

Furthermore, to enhance cell type resolution, we also conducted further subtype annotations and classifications for six major cell types, including: T cells, natural killer cells, myeloid cells, endothelial cells, B cells, and fibroblasts with sophisticated reference data (6, 26-30) (Table S1). For the first four cell types, a larger-scale single-cell atlas was used as the reference dataset, combined with the SingleR(31) and scibetR(32) tool for automatic refined annotation process. Then, the re-annotated results were compared with the cell labels in the original literature; for the latter two cell types, due to the lack of large reference datasets available, typical subtype markers were mainly used, combined with SCINA(33) for automatic annotation process. Similarly, the re-annotated results were compared with the labels in the original literature.

Additionally, we utilized cross-tissue data integration and clustering to analyze cell types in the TME. Firstly, we integrated single-cell data from various organs to classify and analyze cell subtypes with Seurat, which led to the identification of tumor-specific subclusters. Secondly, by utilizing Seurat’s CCA algorithm, we combined and clustered multiple datasets within the same gastrointestinal tumor, revealing the differences in gene expression patterns of the same cell type across normal, paracancerous, and tumor tissues.

### Cell ontology tree construction and markers curation

Following the above comprehensive annotation for digestive single-cells, a standardized ontology for cell types was constructed to structurally describe these cell types in the scRNA-seq dataset of PreDigs data. This ontology tree was based on manual curation and derived from the CL reference database. The complete list of digestive cell types was obtained and matched to the existing CL hierarchy using standardized nomenclature. Subsequently, the corresponding CL numbers and parent node cell types for each cell type were compiled and organized. A unified and structured cell ontology tree was then constructed, mainly categorizing cells into five groups: immune cells, stromal cells, epithelial cells, secretory cells, and others. The text descriptions corresponding to these cell types were also documented and stored uniformly in the database.

In order to construct a comprehensive cell-type marker set for human digestive organs to profile digestive cellular characteristic well, we performed manual curation from current sources of databases. We surveyed well-known cell marker database of Cell Marker2.0(13) and CellMatch(16) and collected canonical markers which all genes were solidly validated by experiment.

### Approach for deriving three context-specific cell markers

To uncover the expression differences of the same cell type across different tissue types, we employed the Seurat package to conduct three distinct differential gene expression analyses for specific cell types under various conditions, including ‘Cell Markers’, ‘Subtype Markers’, and ‘TPN Markers’. Firstly, significantly up-regulated differentially expressed genes (DEGs) in the specific cell type, as compared to other cell types, were identified within each tailored dataset (‘Cell Markers’). The heterogeneity of cell type-specific DEGs across different tissues was elucidated through the comparison of these DEG lists. Subsequently, all datasets across digestive organs containing the specific cell type were integrated to construct a large-scale cell integration dataset. On this integrated dataset, the expression differences between subtypes within the same cell type were analyzed (‘Subtype Markers’). Finally, to meet the research needs specific to a particular digestive organ, relevant datasets were consolidated, and an in-depth analysis of the gene expression differences for the specific cell type across different tissue types (N, P, T) was carried out after eliminating batch effects (‘TPN Markers’).

### Visualization for scRNA-seq data analysis

Except elementary visualization of cell type component statistic and the DEG analysis, UMAPPlot and FeaturePlot function in Seurat were employed for cell clustering visualization was and feature expression. Cell–cell interaction networks were constructed by CellChat (34). Protein-Protein Interaction Networks were constructed by STRINGdb(35) v 2.10.1. KEGG enrichment analysis on differentially expressed genes was performed for each specific cell type using R package clusterProfiler (36) v3.14.3. Survival analysis was performed by R package survival v3.2.7. Static figures in the database were created by ggplot2 v3.3.3.

### Database construction

PreDigs web portal was constructed by standard database development techniques. HTML5 and CSS were used for front-end pages display and MySQL was served as a container for back-end data storage. Echarts and Highcharts were adopted for building interactive graphs. The following browsers were recommended for better compatibility: Google Chrome, Firefox (v64.0 and up), or IE (version 11.0 and up). The whole bioinformatic analysis results in PreDigs were generated by in-house R scripts.

## Data availability

All the data showed in PreDigs can be downloaded from our website at https://www.biosino.org/predigs/. Users are able to access any data and function without registration or login. We also provide a stand-alone R package PreDigsR for the cell type annotation tool and made it available on GitHub. The link to the repository is https://github.com/BioMedBigDataCenter/predigs.

## CRediT author statement

Guoqing Zhang and Liyun Yuan: Conceptualization, Projection administration, Writing-original draft. Jiayue Meng: Investigation, Formal analysis, Data Curation, Writing-original draft. Mengyao Han: Data Curation. Yuwei Huang: Resources. Liang Li and Xiaoyi Chen: Investigation. Yuanhu Ju and Daqing Lv: Software.

## Acknowledgements

This work was supported by grants from the National Major Scientific Instrument and Equipment Development Project of NSFC (81827901), the Strategic Priority Research Program of the Chinese Academy of Sciences (XDB38030100), National Key R&D Program of China (2021YFF0703702), Phase II External Project of Ningbo Institute of Life and Health Industry of University of Chinese Academy of Sciences (2020YJY0217) and the Science and Technology Project of Yunnan Province (202103AQ100002). The work was also supported by Shandong Academician Workstation Program (GP ZHAO) and Shanghai Municipal Science and Technology Major Project. The authors acknowledge the authors from published studies to share their single-cell RNA-seq data on human digestive samples.

## Supplementary material

Figure S1 Data processing and construction workflow of PreDigs

Figure S2 The screenshot of the “Subtype Markers” page in PreDigs

Figure S3 The screenshot of the “TPN Markers” page in PreDigs

Table S1 Cell subtype annotation strategies and reference data types

## References

1. Siegel RL, Giaquinto AN, Jemal A. Cancer statistics, 2024. CA: A Cancer Journal for Clinicians 2024;74:12–49.

2. Zhang Z, Wang Z-X, Chen Y-X, Wu H-X, Yin L, Zhao Q, Luo H-Y, et al. Integrated analysis of single-cell and bulk RNA sequencing data reveals a pan-cancer stemness signature predicting immunotherapy response. Genome Medicine 2022;14.

3. Maman S, Witz IP. A history of exploring cancer in context. Nature Reviews Cancer 2018;18:359–376.

4. Wang Y, Navin Nicholas E. Advances and Applications of Single-Cell Sequencing Technologies. Molecular Cell 2015;58:598–609.

5. Kester L, van Oudenaarden A. Single-Cell Transcriptomics Meets Lineage Tracing. Cell Stem Cell 2018;23:166–179.

6. Cheng S, Li Z, Gao R, Xing B, Gao Y, Yang Y, Qin S, et al. A pan-cancer single-cell transcriptional atlas of tumor infiltrating myeloid cells. Cell 2021;184:792-809.e723.

7. Ono H, Arai Y, Furukawa E, Narushima D, Matsuura T, Nakamura H, Shiokawa D, et al. Single-cell DNA and RNA sequencing reveals the dynamics of intra-tumor heterogeneity in a colorectal cancer model. BMC Biology 2021;19.

8. Croft W, Evans RPT, Pearce H, Elshafie M, Griffiths EA, Moss P. The single cell transcriptional landscape of esophageal adenocarcinoma and its modulation by neoadjuvant chemotherapy. Molecular Cancer 2022;21.

9. Ferrall-Fairbanks MC, Ball M, Padron E, Altrock PM. Leveraging Single-Cell RNA Sequencing Experiments to Model Intratumor Heterogeneity. JCO Clinical Cancer Informatics 2019:1–10.

10. Gao R, Bai S, Henderson YC, Lin Y, Schalck A, Yan Y, Kumar T, et al. Delineating copy number and clonal substructure in human tumors from single-cell transcriptomes. Nature Biotechnology 2021;39:599–608.

11. Diehl AD, Meehan TF, Bradford YM, Brush MH, Dahdul WM, Dougall DS, He Y, et al. The Cell Ontology 2016: enhanced content, modularization, and ontology interoperability. Journal of Biomedical Semantics 2016;7.

12. Zhang X, Lan Y, Xu J, Quan F, Zhao E, Deng C, Luo T, et al. CellMarker: a manually curated resource of cell markers in human and mouse. Nucleic Acids Research 2019;47:D721–D728.

13. Hu C, Li T, Xu Y, Zhang X, Li F, Bai J, Chen J, et al. CellMarker 2.0: an updated database of manually curated cell markers in human/mouse and web tools based on scRNA-seq data. Nucleic Acids Research 2023;51:D870–D876.

14. Hatano A, Chiba H, Moesa HA, Taniguchi T, Nagaie S, Yamanegi K, Takai-Igarashi T, et al. CELLPEDIA: a repository for human cell information for cell studies and differentiation analyses. Database 2011;2011:bar046–bar046.

15. Franzén O, Gan L-M, Björkegren JLM. PanglaoDB: a web server for exploration of mouse and human single-cell RNA sequencing data. Database 2019;2019.

16. Shao X, Liao J, Lu X, Xue R, Ai N, Fan X. scCATCH: Automatic Annotation on Cell Types of Clusters from Single-Cell RNA Sequencing Data. iScience 2020;23.

17. Choi J-H, In Kim H, Woo HG. scTyper: a comprehensive pipeline for the cell typing analysis of single-cell RNA-seq data. BMC Bioinformatics 2020;21.

18. Stachelscheid H, Seltmann S, Lekschas F, Fontaine J-F, Mah N, Neves M, Andrade-Navarro MA, et al. CellFinder: a cell data repository. Nucleic Acids Research 2014;42:D950–D958.

19. Jiang S, Qian Q, Zhu T, Zong W, Shang Y, Jin T, Zhang Y, et al. Cell Taxonomy: a curated repository of cell types with multifaceted characterization. Nucleic Acids Research 2023;51:D853–D860.

20. Zhang Y, Sun H, Zhang W, Fu T, Huang S, Mou M, Zhang J, et al. CellSTAR: a comprehensive resource for single-cell transcriptomic annotation. Nucleic Acids Research 2024;52:D859–D870.

21. Han X, Zhou Z, Fei L, Sun H, Wang R, Chen Y, Chen H, et al. Construction of a human cell landscape at single-cell level. Nature 2020;581:303–309.

22. McGinnis CS, Murrow LM, Gartner ZJ. DoubletFinder: Doublet Detection in Single-Cell RNA Sequencing Data Using Artificial Nearest Neighbors. Cell Syst 2019;8:329–337 e324.

23. Hao Y, Hao S, Andersen-Nissen E, Mauck WM, Zheng S, Butler A, Lee MJ, et al. Integrated analysis of multimodal single-cell data. Cell 2021;184:3573-3587.e3529.

24. Korsunsky I, Millard N, Fan J, Slowikowski K, Zhang F, Wei K, Baglaenko Y, et al. Fast, sensitive and accurate integration of single-cell data with Harmony. Nature Methods 2019;16:1289–1296.

25. Zhang AW, O’Flanagan C, Chavez EA, Lim JLP, Ceglia N, McPherson A, Wiens M, et al. Probabilistic cell-type assignment of single-cell RNA-seq for tumor microenvironment profiling. Nature Methods 2019;16:1007–1015.

26. Chu Y, Dai E, Li Y, Han G, Pei G, Ingram DR, Thakkar K, et al. Pan-cancer T cell atlas links a cellular stress response state to immunotherapy resistance. Nature Medicine 2023;29:1550–1562.

27. Tang F, Li J, Qi L, Liu D, Bo Y, Qin S, Miao Y, et al. A pan-cancer single-cell panorama of human natural killer cells. Cell 2023;186:4235-4251.e4220.

28. Xia J, Xie Z, Niu G, Lu Z, Wang Z, Xing Y, Ren J, et al. Single-cell landscape and clinical outcomes of infiltrating B cells in colorectal cancer. Immunology 2022;168:135–151.

29. Lavie D, Ben-Shmuel A, Erez N, Scherz-Shouval R. Cancer-associated fibroblasts in the single-cell era. Nature Cancer 2022;3:793–807.

30. Dai J, Xi X, Liu Z, Wu W, Zhu S, Zhang X, Huang Y, et al. Single-cell sequencing of multi-region resolves geospatial architecture and therapeutic target of endothelial cells in esophageal squamous cell carcinoma. Clinical and Translational Medicine 2023;13.

31. Aran D, Looney AP, Liu L, Wu E, Fong V, Hsu A, Chak S, et al. Reference-based analysis of lung single-cell sequencing reveals a transitional profibrotic macrophage. Nature Immunology 2019;20:163–172.

32. Li C, Liu B, Kang B, Liu Z, Liu Y, Chen C, Ren X, et al. SciBet as a portable and fast single cell type identifier. Nature Communications 2020;11.

33. Zhang Z, Luo D, Zhong X, Choi JH, Ma Y, Wang S, Mahrt E, et al. SCINA: A Semi-Supervised Subtyping Algorithm of Single Cells and Bulk Samples. Genes 2019;10.

34. Jin S, Guerrero-Juarez CF, Zhang L, Chang I, Ramos R, Kuan C-H, Myung P, et al. Inference and analysis of cell-cell communication using CellChat. Nature Communications 2021;12.

35. Szklarczyk D, Gable AL, Nastou KC, Lyon D, Kirsch R, Pyysalo S, Doncheva NT, et al. The STRING database in 2021: customizable protein–protein networks, and functional characterization of user-uploaded gene/measurement sets. Nucleic Acids Research 2021;49:D605–D612.

36. Wu T, Hu E, Xu S, Chen M, Guo P, Dai Z, Feng T, et al. clusterProfiler 4.0: A universal enrichment tool for interpreting omics data. The Innovation 2021;2.

37. Mayer S, Milo T, Isaacson A, Halperin C, Miyara S, Stein Y, Lior C, et al. The tumor microenvironment shows a hierarchy of cell-cell interactions dominated by fibroblasts. Nature Communications 2023;14.

