## Supplementary material for "PreDigs: a Database of Context-specific Cell-type Markers and Precise cell subtypes for Digestive Cell Annotation"

### Supplemental materials

Figure S1 Data processing and construction workflow of PreDigs

Figure S2 The screenshot of the "Subtype Markers" page in PreDigs

Figure S3 The screenshot of the "TPN Markers" page in PreDigs

Table S1 Cell subtype annotation strategies and reference data types

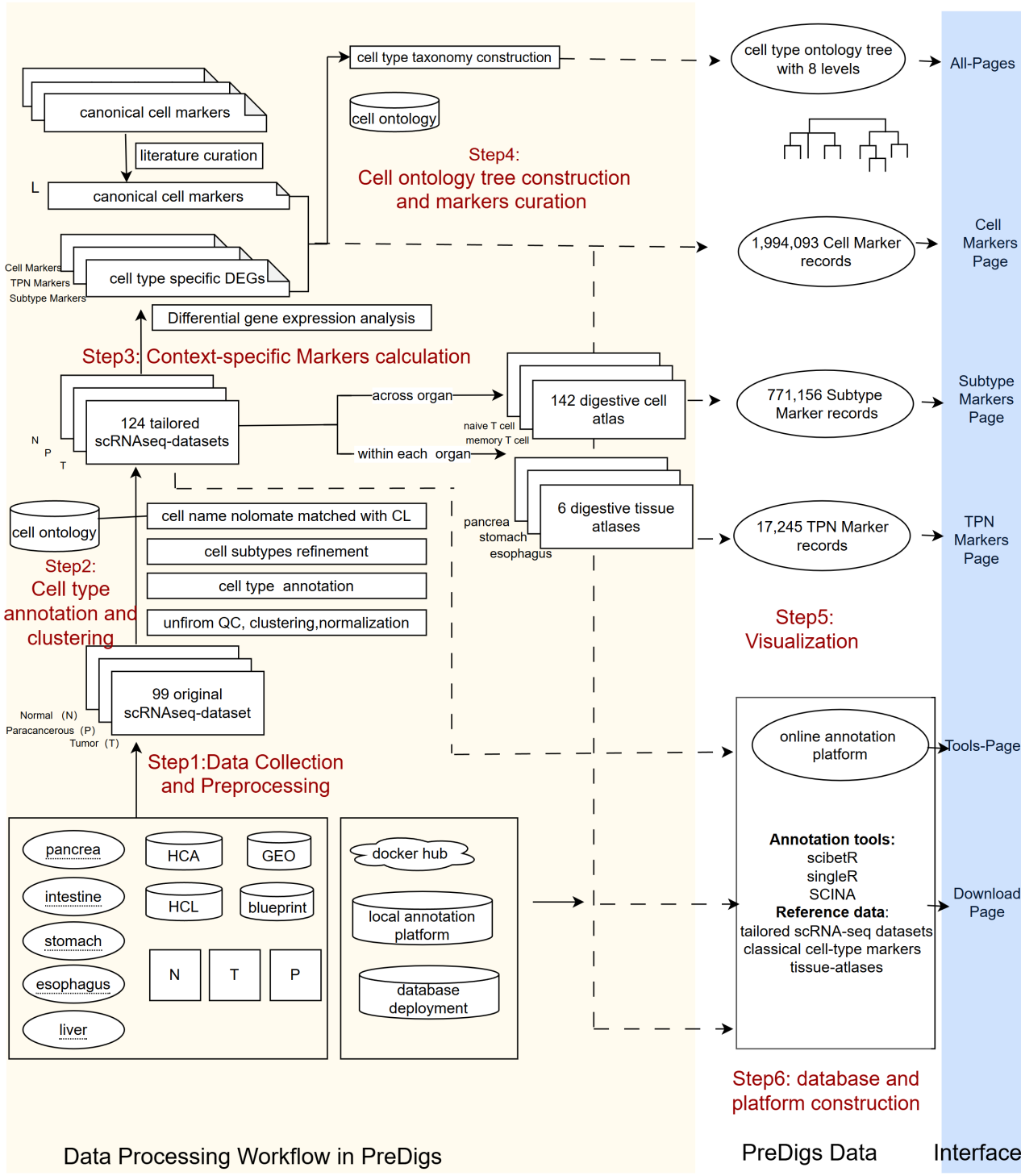

**FigureS1** Data processing and database construction workflow of PreDigs

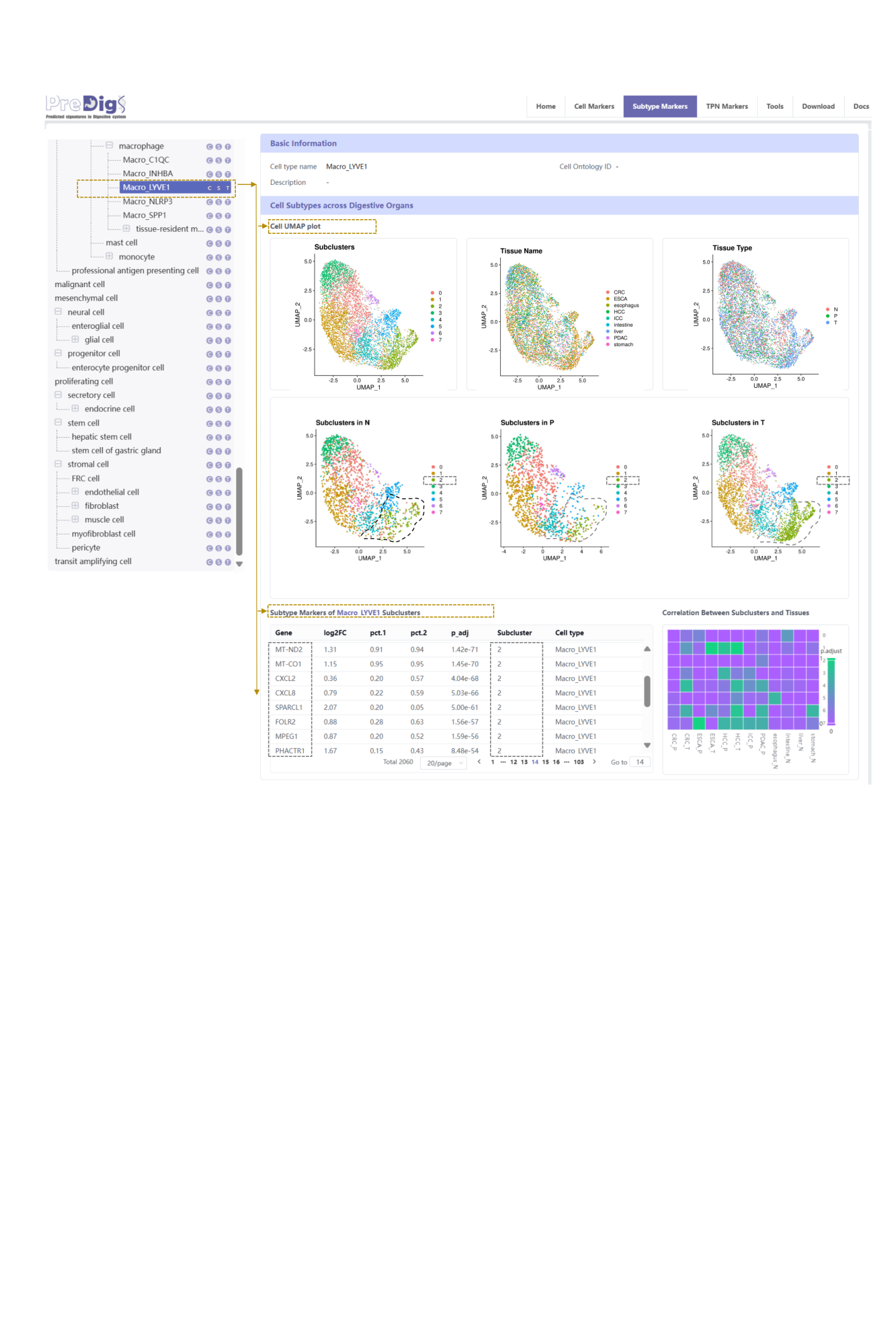

**Figure S2** The screenshot of the "Subtype Markers" page in PreDigs

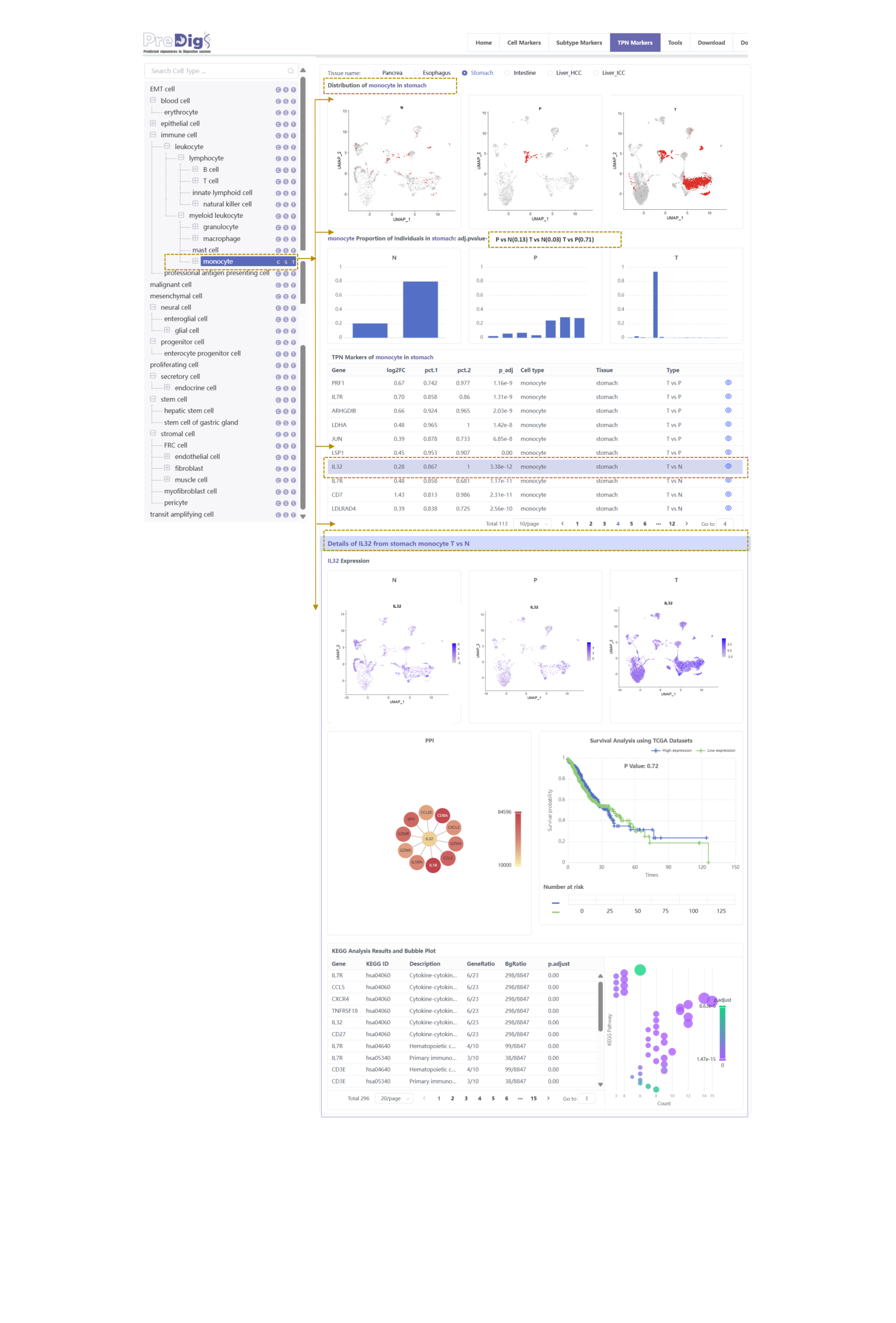

**Figure S3** The screenshot of the "TPN Markers" page in PreDigs

**Table S1**　Cell Subtype Annotation Strategies and Reference Data Types

| Cell Type | Cell Subtype | Reference Data | Reference |
| --- | --- | --- | --- |
| T cell | Teff, Th17,Tfh,  CTL,Treg,Tem,  Tcm,Tn,Tex,NKT,  gamma-delta T | gene expression matrix | Chu Y,et al.Nat Med.2023  (37248301) |
| natural killer cell | CD56brightCDlowNK,CD56dimCD16hiNK | gene expression matrix | Tang F,et al.Cell.2023  (37607536) |
| myeloid leukocyte | cDC,  neutrophil,  mast cell,  classical monocyte,  non-classical monocyte | gene expression matrix | Cheng S,et al.Cell.2021  (33545035) |
| endothelial cell | lymphatic, artery, vein, tip cell, TEC,  stalk cell, capilary | gene expression matrix | Dai J,et al. Clin Transl Med. 2023. (37987158) |
| B cell | plasma, naive B,  memory B | Plasma.cell：  SDC1,MZB1,IGHG1,IGHA1,IGKC  Naive.B.cell：MS4A1,IGHD,FCER2,TCL1A,IL4R  Memory.B.cell：MS4A1,CD27,AIM2,TNFRSF13B | Xia J,et al.Immunology.2023  (36082430) |
| fibroblast | myofibroblastic CAF (myoCAF)  immune regulatory& inflamatory(iCAF) | myoCAF：  COL1A1,COL10A1,COL4A1,MMP3,IL4,IL13,TGFB1,ACTG2,ACTA2,FAP,PDPN  iCAF：  IL6,IL11,IL8,LIF,CSF2,CXCL1,CXCL12,CXCL14,CCL2,CCL8,CFD,C1QC,C1QA,C1QC,HLA-DRA | Lavie D,et al.Nat Cancer.2022  (35883004) |
